# Nelfinavir was predicted to be a potential inhibitor of 2019-nCov main protease by an integrative approach combining homology modelling, molecular docking and binding free energy calculation

**DOI:** 10.1101/2020.01.27.921627

**Authors:** Zhijian Xu, Cheng Peng, Yulong Shi, Zhengdan Zhu, Kaijie Mu, Xiaoyu Wang, Weiliang Zhu

## Abstract

2019-nCov has caused more than 80 deaths as of 27 January 2020 in China, and infection cases have been reported in more than 10 countries. However, there is no approved drug to treat the disease. 2019-nCov M^pro^ is a potential drug target to combat the virus. We built homology models based on SARS M^pro^ structures, and docked 1903 small molecule drugs to the models. Based on the docking score and the 3D similarity of the binding mode to the known M^pro^ ligands, 4 drugs were selected for binding free energy calculations. Both MM/GBSA and SIE methods voted for nelfinavir, with the binding free energy of −24.69±0.52 kcal/mol and −9.42±0.04 kcal/mol, respectively. Therefore, we suggested that nelfinavir might be a potential inhibitor against 2019-nCov M^pro^.

## 1. Introduction

In December 2019, cluster of patients with pneumonia were reported in Wuhan, Hubei Province, China.^1, 2^ Shortly, a new coronavirus, temporally named 2019-nCov, was identified to be the cause of the disease, which is the seventh member of the family betacoronavirus.^1^ More than 2,700 infection cases and 80 deaths were reported as of 27 January 2020 from China. In addition, infection cases have been reported from Thailand, Australia, Malaysia, Singapore, France, Japan, South Korea, the United States, Vietnam, Canada and Nepal, indicating that the disease is a potential threat to the global health.^3^ Sadly, the number is still growing rapidly and no drug has been approved to be effective. Therefore, it is urgent to discover and develop drugs to cure the disease.

Based on its function, the main protease (M^pro^) or chymotrypsin-like protease (3CL^pro^)^4^ is suggested to be a potential drug target to combat 2019-nCov, which is highly conservable among coronaviruses. Sequence alignment revealed that the M^pro^ of 2019-nCov shares 96% similarity with that of SARS (severe acute respiratory syndrome) (Figure 1). Studies for identifying the inhibitors of 2019-nCov M^pro^ were quickly performed for discovering and developing drugs against the disease. For example, Hualiang Jiang and collaborators identified 30 drugs and compounds as the M^pro^ inhibitors via protein modelling and virtual screening, which is a rapid progress in the way to cope with the crisis. In addition, one of the 30 drugs/compounds, remdesivir, was also suggested to be potential inhibitor against 2019-nCov by Liu et al,^5^ who also suggested 3 other possible inhibitors.

**Figure 1.**
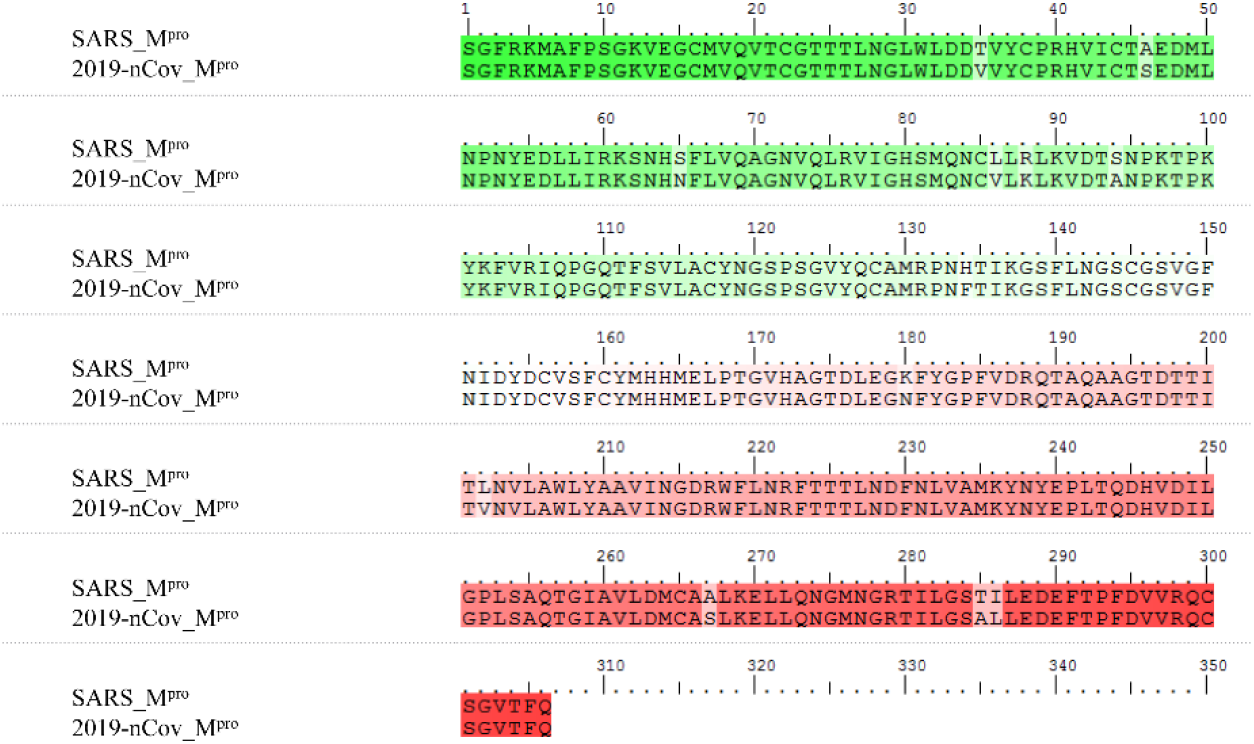
Sequence alignment of 2019-nCov M^pro^ and SARS M^pro^.

For finding more potential drugs as inhibitors of the protein, we modelled the 2019-nCov M^pro^ structures using SARS M^pro^ (PDB ID: 2GTB) as template and docked 1903 approved drugs against the model in this study. Fifteen drugs were selected based on the docking score and three dimensional (3D) similarity to available M^pro^ inhibitors against other coronavirus. For validation, we modelled 10 additional new models of 2019-nCov M^pro^ and docked the 15 drugs against the new models, which revealed that 6 drugs (nelfinavir, praziquantel, pitavastatin, perampanel, eszopiclone, and zopiclone) have good binding modes with the new models. Binding free energy calculation were then performed for 4 of the 6 drugs using MM/GBSA and SIE methods. Nelfinavir was identified as the best one with predicted binding free energies of −24.69±0.52 kcal/mol by MM/GBSA and −9.42±0.04 kcal/mol by SIE, respectively. Taking into account its high 3D similarity of binding mode to the known M^pro^ inhibitor, we proposed that nelfinavir should be a potential inhibitor against 2019-nCov M^pro^.

## 2. Materials and methods

### 2.1 Homology modelling

43 M^pro^ complexes with ligands were downloaded from protein data bank^6^ (PDB IDs: 1WOF, 2A5I, 2A5K, 2ALV, 2AMD, 2GTB, 2GX4, 2GZ7, 2GZ8, 2OP9, 2QIQ, 2V6N, 2ZU4, 2ZU5, 3SN8, 3SND, 3SZN, 3TIT, 3TIU, 3TNS, 3TNT, 3V3M, 4F49, 4MDS, 4TWW, 4TWY, 4WY3, 4YLU, 4YOG, 4YOI, 4YOJ, 4ZRO, 5C5N, 5C5O, 5EU8, 5N5O, 5N19, 5NH0, 5WKJ, 5WKK, 5WKL, 5WKM,6FV1) and aligned to 2GTB in PyMOL.^7^ 11 complexes (PDB IDs: 2A5K, 2GTB, 2GX4, 3SND, 3TNS, 3V3M, 4F49, 4YLU, 5NH0, 5WKJ, 5WKM) were served as templates to build 11 2019-nCov M^pro^ models in SWISS-MODEL server by “user template” mode.^8^

### 2.2 Approved Drugs

1905 approved small molecule drugs with 3D coordinates were downloaded from DrugBank release version 5.1.5,^9^ while 1903 drugs could be converted to pdbqt format by prepare_ligand4.py script in MGLToos version 1.5.6.^10^

### 2.3 Molecular Docking

1903 approved drugs in pdbqt format were docked to 2019-nCov M^pro^ model (template: 2GTB) by smina,^11^ which is a fork of AutoDock Vina^12^ with improving scoring and minimization. The hydrogens were added to 2019-nCov M^pro^ model by pdb2pqr (--ff=amber --ffout=amber --chain --with-ph=7).^13^ Then the model was converted to pdbqt format by prepare_receptor4.py script in MGLToos version 1.5.6.^10^ The ligand in 2GTB was used to define the grid and the buffer space was set to 6.0 Å (autobox_add). The random seed was explicitly set to 0 (seed). The exhaustiveness of the global search was set to 32 (exhaustiveness) and at most 1 binding mode was generated for each drug (num_modes). MolShaCS, which utilized Gaussian-based description of molecular shape and charge distribution, was used to calculate the 3D similarities between approved drugs and available M^pro^ inhibitors.^14^

### 2.4 Molecular dynamics simulation

Each simulation system was immersed in a cubic box of TIP3P water that was extended by 9 Å from the solute, with a rational number of counter ions of Na^+^ or Cl^−^ to neutralize the system. General Amber force field (GAFF)^15^ and Amber ff03 force field^16^ were used to parameterize the ligand and protein, respectively. 10,000 steps of minimization with constraints (10 kcal/mol/Å^2^) on heavy atoms of complex, including 5,000 steps of steepest descent minimization and 5,000 steps of conjugate gradient minimization, was used to optimize each system. Then each system was heated to 300 K within 0.2 ns followed by 0.1 ns equilibration in NPT ensemble. Finally, 5 ns MD simulation on each system at 300 K was performed. The minimization, heating and equilibrium are performed with *sander* program in Amber16. The 5 ns production run was performed with *pmemd.cuda*.

### 2.5 Binding free energy calculation

Based on the 5 ns MD simulation trajectory, bind-free energy (Δ*G*) was calculated with MM/GBSA^17, 18^ and SIE^19^ approaches. In the MM/GBSA, the Δ*G* was calculated according to equation (1),

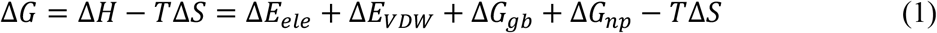

where Δ*E*_ele_ and Δ*E*_VDW_ refer to electrostatic and van der Waals energy terms, respectively. Δ*G*_gb_ and Δ*G*_np_ refer to polar and non-polar solvation free energies, respectively. Conformational entropy (*T*Δ*S*) was calculated by *nmode* module in Amber16. The dielectric constants for solvent and solute were set to 80.0 and 1.0, respectively, and OBC solvation model (igb = 5 and PBradii = 5)^20^ was used in this study. Other parameters are set to default values.

In the SIE, the Δ*G* was calculated based on equation (2),

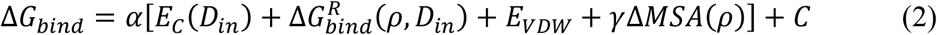

where *E*_C_ and *E*_VDW_ refer to the sum of intermolecular Coulomb and van der Waals interaction energies, respectively. 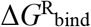 and Δ*MSA* refer to the changes of reaction field energy and molecular surface area upon ligand binding, respectively. Default values of the global proportionality coefficient (*α* = 0.1048), the solute interior dielectric constant (*D_in_* = 2.25), the van der Waals radii linear scaling coefficient (*ρ* = 1.1), the molecular surface area coefficient (*γ* = 0.0129 kcal/mol/Å^2^), and the constant (*C* = −2.89 kcal/mol) are used in this study.

## 3. Results and discussion

### 3.1 Docking results of 1903 approved drugs against M^pro^ models

1903 approved small molecule drugs were docked to a M^pro^ homology model built by SWISS-MODEL using 2GTB as template. There are 672 drugs with the docking score better than −7.0 kcal/mol. The binding modes are essential to the activities. If a compound shares a similar binding mode to a known ligand, it is more likely to have a similar activity to the ligand. Therefore, we calculated the 3D similarities of the binding mode of the docked drugs to the available binders against M^pro^. 39 binders in 44 complexes were chosen as the references to calculate the 3D similarities (Table 1). 159 drugs have a 3D similarity larger than 65.0% with at least one known M^pro^ binder.

**Table 1.**
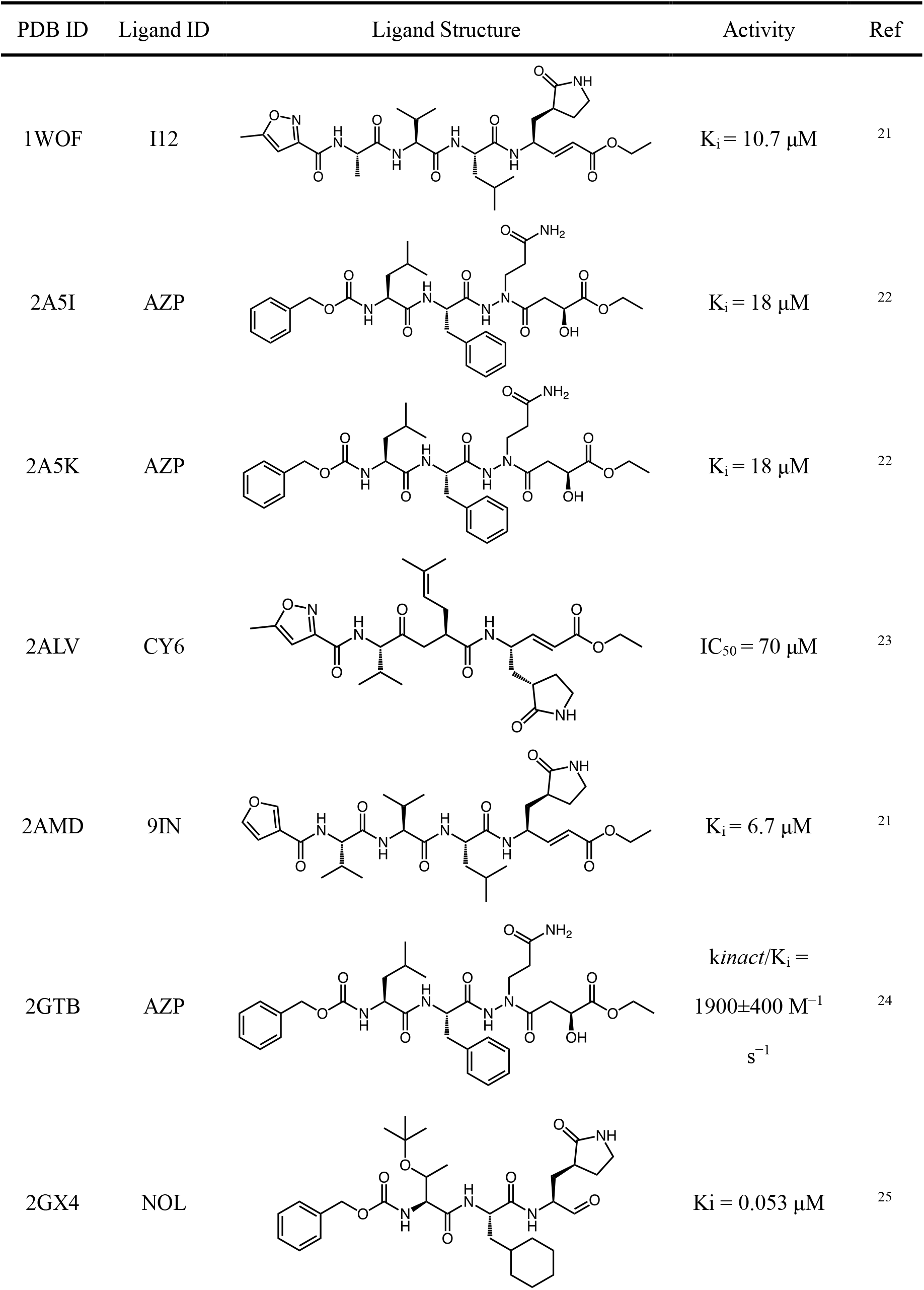
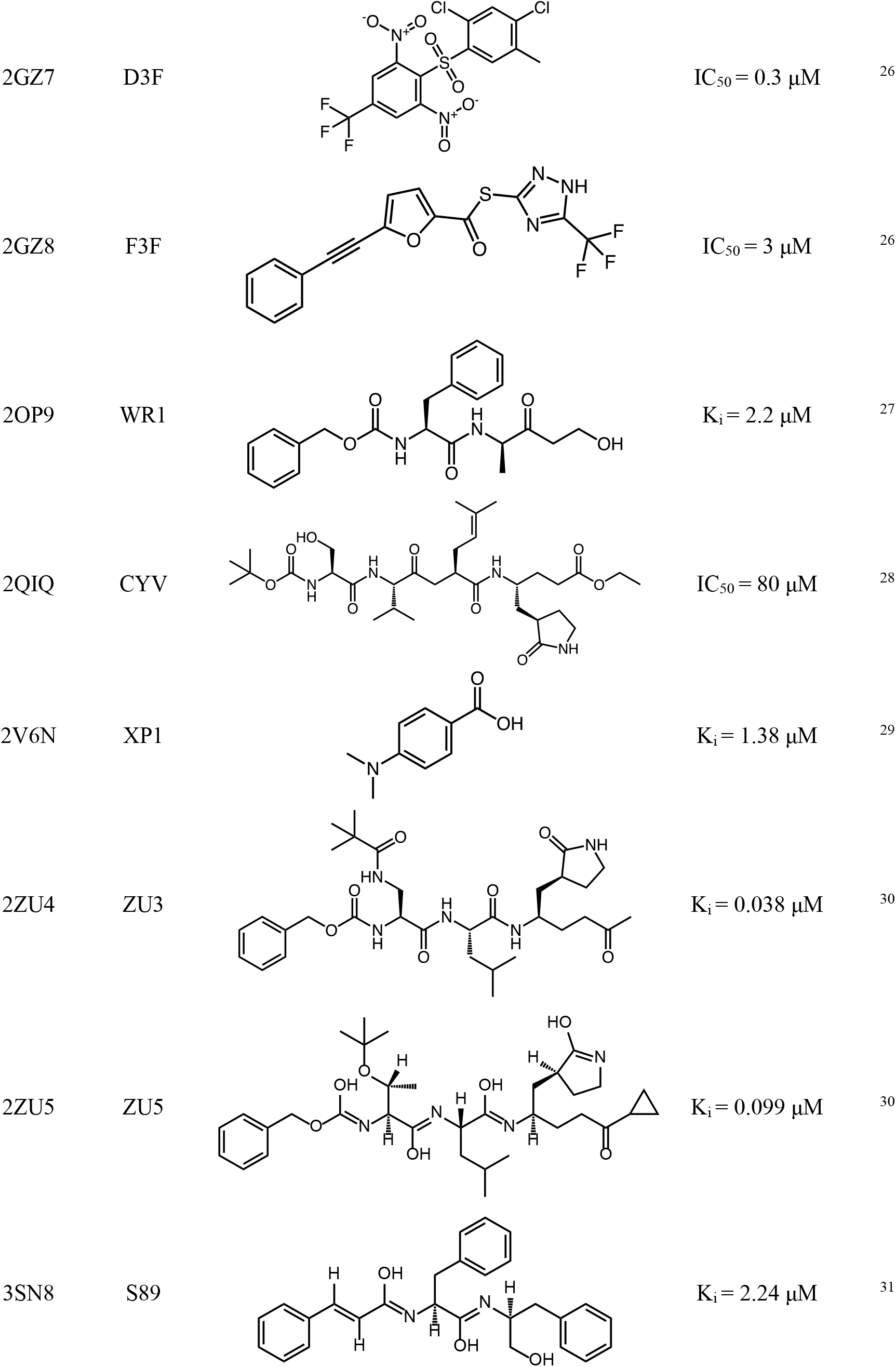
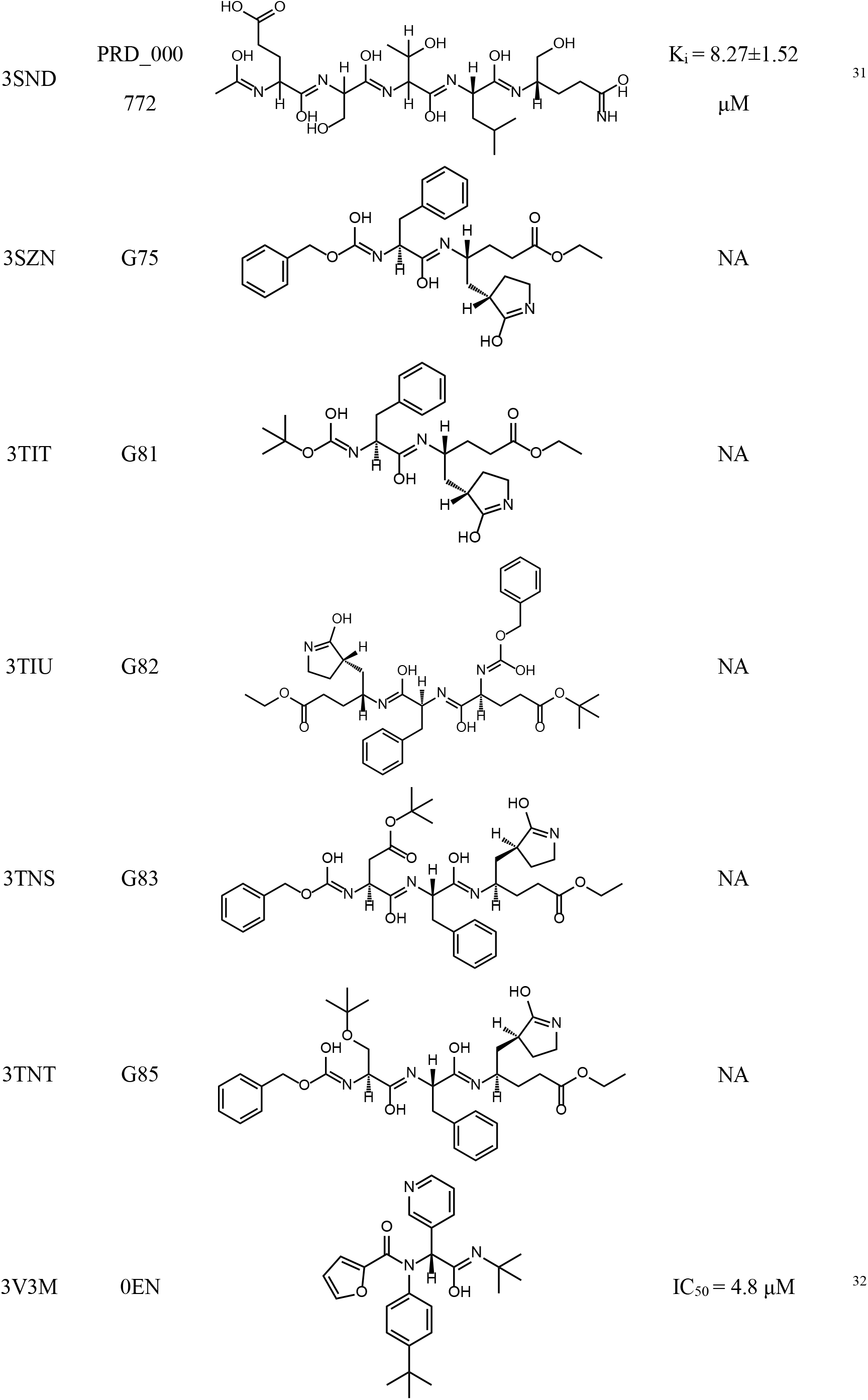
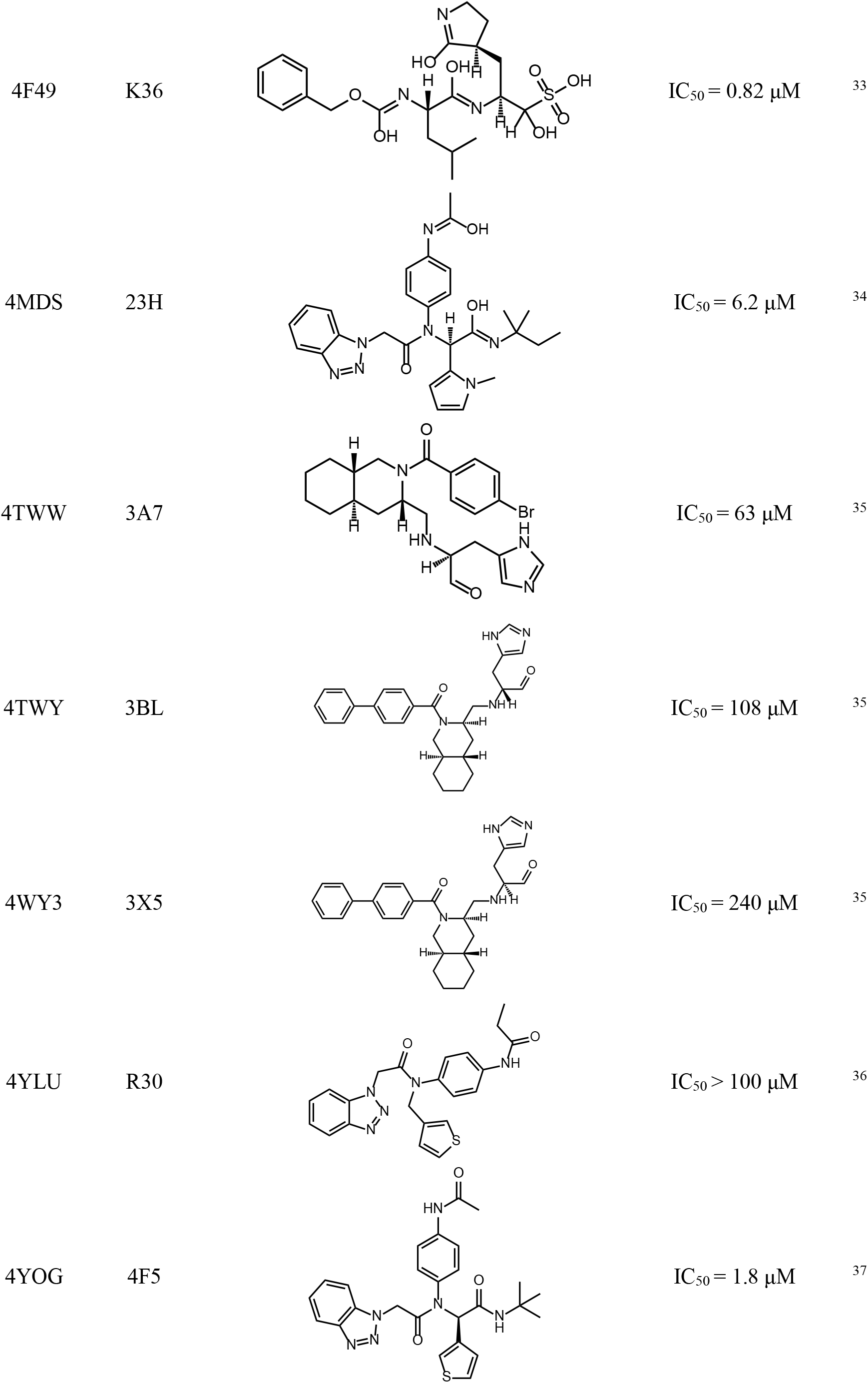
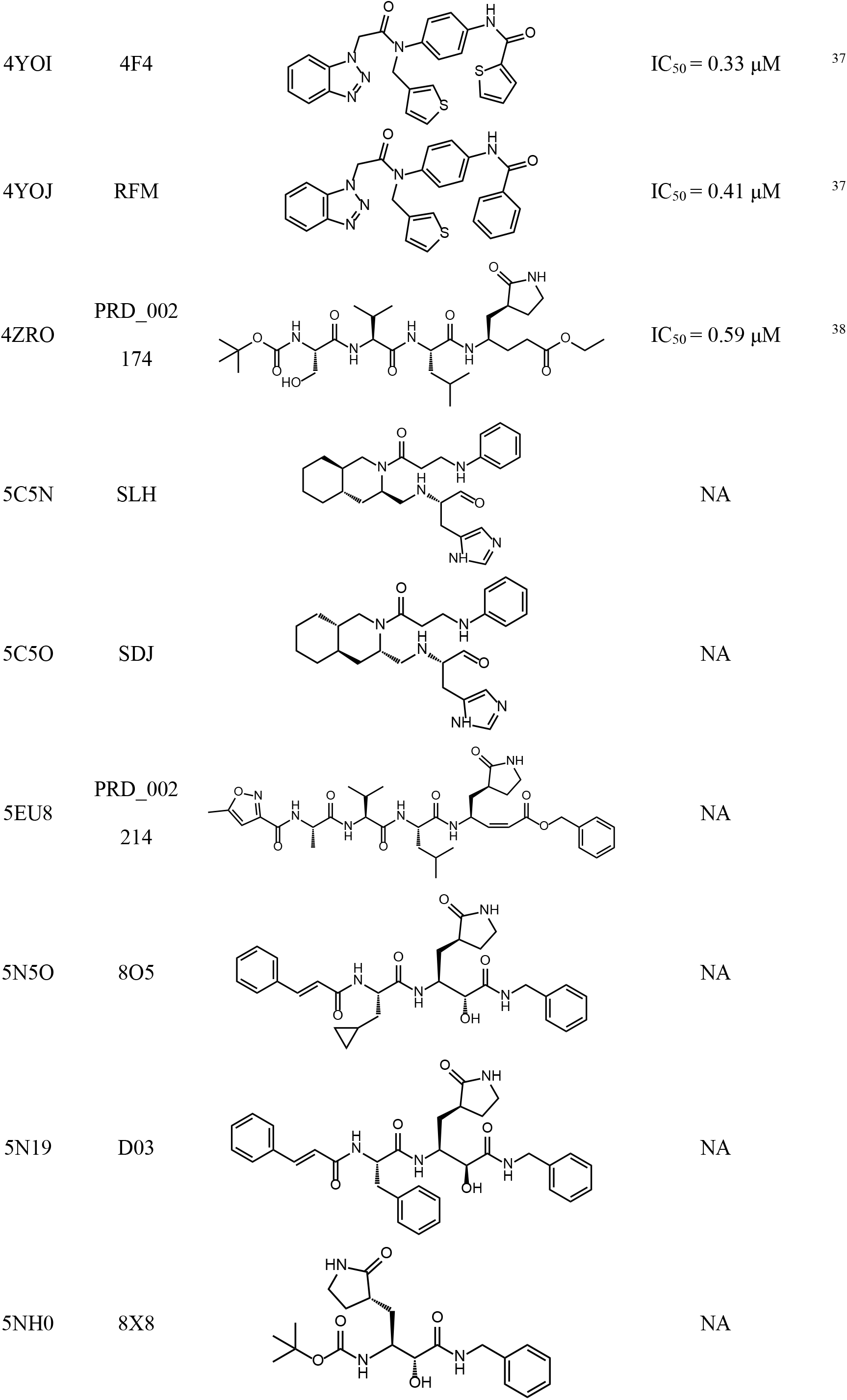
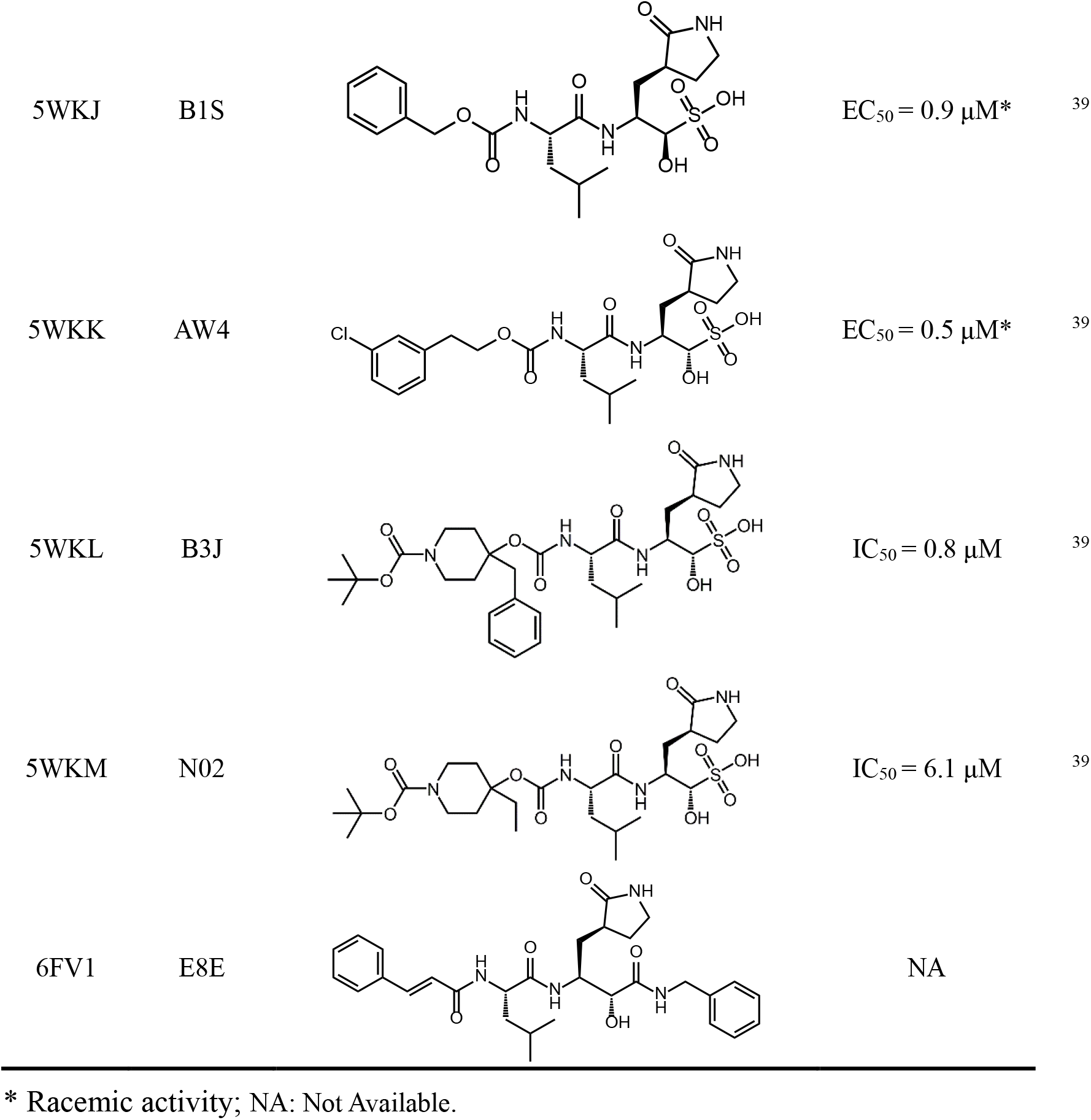
Known M^pro^ binders as references to calculate the 3D similarity of binding modes.

| PDB ID | Ligand ID | Ligand Structure | Activity | Ref |
| --- | --- | --- | --- | --- |
| 1WOF | I12 | | $K_i = 10.7 \mu\text{M}$ | 21 |
| 2A5I | AZP | | $K_i = 18 \mu\text{M}$ | 22 |
| 2A5K | AZP | | $K_i = 18 \mu\text{M}$ | 22 |
| 2ALV | CY6 | | $\text{IC}_{50} = 70 \mu\text{M}$ | 23 |
| 2AMD | 9IN | | $K_i = 6.7 \mu\text{M}$ | 21 |
| 2GTB | AZP | | $k_{\text{inact}}/K_i = 1900 \pm 400 \text{ M}^{-1} \text{ s}^{-1}$ | 24 |
| 2GX4 | NOL | | $K_i = 0.053 \mu\text{M}$ | 25 |
| 2GZ7 | D3F | | $IC_{50} = 0.3 \mu M$ | 26 |
| 2GZ8 | F3F | | $IC_{50} = 3 \mu M$ | 26 |
| 2OP9 | WR1 | | $K_i = 2.2 \mu M$ | 27 |
| 2QIQ | CYV | | $IC_{50} = 80 \mu M$ | 28 |
| 2V6N | XP1 | | $K_i = 1.38 \mu M$ | 29 |
| 2ZU4 | ZU3 | | $K_i = 0.038 \mu M$ | 30 |
| 2ZU5 | ZU5 | | $K_i = 0.099 \mu M$ | 30 |
| 3SN8 | S89 | | $K_i = 2.24 \mu M$ | 31 |
| 3SND | PRD_000<br>772 | | $K_i = 8.27 \pm 1.52$<br>$\mu\text{M}$ | 31 |
| 3SZN | G75 |  | NA |  |
| 3TIT | G81 |  | NA |  |
| 3TIU | G82 |  | NA |  |
| 3TNS | G83 |  | NA |  |
| 3TNT | G85 |  | NA |  |
| 3V3M | 0EN | | $\text{IC}_{50} = 4.8 \mu\text{M}$ | 32 |
| 4F49 | K36 | | $IC_{50} = 0.82 \mu M$ | 33 |
| 4MDS | 23H | | $IC_{50} = 6.2 \mu M$ | 34 |
| 4TWW | 3A7 | | $IC_{50} = 63 \mu M$ | 35 |
| 4TWY | 3BL | | $IC_{50} = 108 \mu M$ | 35 |
| 4WY3 | 3X5 | | $IC_{50} = 240 \mu M$ | 35 |
| 4YLU | R30 | | $IC_{50} > 100 \mu M$ | 36 |
| 4YOG | 4F5 | | $IC_{50} = 1.8 \mu M$ | 37 |
| 4YOI | 4F4 | | $IC_{50} = 0.33 \mu M$ | 37 |
| 4YOJ | RFM | | $IC_{50} = 0.41 \mu M$ | 37 |
| 4ZRO | PRD_002<br>174 | | $IC_{50} = 0.59 \mu M$ | 38 |
| 5C5N | SLH |  | NA |  |
| 5C5O | SDJ |  | NA |  |
| 5EU8 | PRD_002<br>214 |  | NA |  |
| 5N5O | 8O5 |  | NA |  |
| 5N19 | D03 |  | NA |  |
| 5NH0 | 8X8 |  | NA |  |
| 5WKJ | B1S |  | EC<sub>50</sub> = 0.9 μM* | 39 |
| 5WKK | AW4 |  | EC<sub>50</sub> = 0.5 μM* | 39 |
| 5WKL | B3J |  | IC<sub>50</sub> = 0.8 μM | 39 |
| 5WKM | N02 |  | IC<sub>50</sub> = 6.1 μM | 39 |
| 6FV1 | E8E |  | NA |  |

\* Racemic activity; NA: Not Available.

After visualizing the docked complexes carefully, we selected 15 drugs (Table 2) for further analysis. There are some differences between the conformations of the protein in the 3D similarity reference and 2GTB model. Therefore, we modelled 10 new homology models using the proteins in 3D similarity reference as templates and we re-docked the 15 drugs to the 10 new homology models. 6 drugs (nelfinavir, pitavastatin, perampanel, praziquantel, zopiclone, and eszopiclone) show good docking scores and binding modes (Table 3). Because eszopiclone and zopiclone was used to treat insomnia in a low dosage which may not be suitable to treat pneumonia, we carried out further binding free energy calculation for the rest 4 drugs.

**Table 2.** 15 drugs selected from 2GTB model.

| DrugBank ID | Docking Score against<br>2GTB model (kcal/mol) | 3D Similarity<br>Reference | 3D<br>Similarity |
| --- | --- | --- | --- |
| DB00328 | -7.16 | 3V3M | 71.3% |
| DB13680 | -7.13 | 3SND | 70.8% |
| DB01165 | -7.31 | 3V3M | 70.2% |
| DB01198 | -7.81 | 5NH0 | 68.9% |
| DB08934 | -8.65 | 2GX4 | 68.6% |
| DB01165 | -7.31 | 5WKM | 68.3% |
| DB08860 | -8.06 | 2GTB | 68.1% |
| DB00402 | -7.75 | 5NH0 | 67.8% |
| DB08883 | -8.17 | 5NH0 | 67.1% |
| DB01288 | -7.96 | 5WKM | 67.1% |
| DB08860 | -8.06 | 2A5K | 66.9% |
| DB00972 | -7.71 | 5NH0 | 66.7% |
| DB00482 | -8.46 | 4YLU | 66.7% |
| DB00220 | -8.91 | 2GX4 | 66.4% |
| DB01058 | -7.90 | 3V3M | 65.7% |
| DB00328 | -7.16 | 4F49 | 65.6% |
| DB08934 | -8.65 | 3TNS | 65.5% |
| DB00904 | -7.45 | 5WKM | 65.4% |
| DB11951 | -8.27 | 4F49 | 65.3% |
| DB11951 | -8.27 | 5WKJ | 65.2% |

**Table 3.** 6 drugs selected from 10 new homology models.

| DrugBank ID | Name | Homology Template | Docking Score (kcal/mol) | 3D Similarity |
| --- | --- | --- | --- | --- |
| DB00220 | nelfinavir | 2GX4 | -9.18 | 70.2% |
| DB08860 | pitavastatin | 2GTB | -8.06 | 68.1% |
| DB08883 | perampanel | 5NH0 | -8.63 | 66.9% |
| DB01058 | praziquantel | 3V3M | -7.38 | 65.8% |
| DB01198 | zopiclone | 5NH0 | -8.46 | 63.5% |
| DB00402 | eszopiclone | 5NH0 | -8.46 | 63.5% |

### 3.2 Binding Free Energy calculated by MM/GBSA and SIE

The 4 docked complex structures, i.e., nelfinavir-2GX4, pitavastatin-2GTB, perampanel-5NH0 and praziquantel-3V3M, were subjected to 5 ns molecular dynamics simulations using Amber 16. To provide insight into their binding mechanisms, the binding free energies were calculated by MM/GBSA and SIE approaches. In results of the MM/GBSA approach, the calculated binding free energies of nelfinavir-2GX4, pitavastatin-2GTB, perampanel-5NH0, and praziquantel-3V3M, are −24.69±0.52, −12.70±0.38, −14.98±0.34, and −6.51±0.21 kcal/mol, respectively (Table 4), which highlight nelfinavir as the most active one. In pitavastatin-2GTB, perampanel-5NH0, and praziquantel-3V3M, the van der Waals interaction (*E*_vdw_) makes a more significant contribution than the electrostatic interaction (*E*_ele_) (Table 4), indicating that van der Waals interaction is the main driving force for the 3 drugs binding. However, the *E*ele interaction of nelfinavir-2GX4 is very strong, suggesting that the electrostatic interaction also play an important role in the binding of nelfinavir. Furthermore, in results of the SIE approach, the rank of the calculated binding free energies for the 4 docking complexes is consistent with that of the MM/GBSA approach (Table 5) that the nelfinavir has the strongest binding free energy (−9.42±0.04 kcal/mol), indicating the reliability of the current binding free energy analysis.

**Table 4.** Components of the Binding Free Energy (kcal/mol) Calculated by MM/GBSA Approach*.

| Energy term | nelfinavir | pitavastatin | perampanel | praziquantel |
| --- | --- | --- | --- | --- |
| $E_{\text{vdw}}$ | $-58.36 \pm 0.32$ | $-42.77 \pm 0.30$ | $-44.33 \pm 0.25$ | $-34.92 \pm 0.32$ |
| $E_{\text{ele}}$ | $-117.95 \pm 0.81$ | $37.95 \pm 0.86$ | $0.05 \pm 0.03$ | $-8.64 \pm 0.35$ |
| $G_{\text{gb}}$ | $134.49 \pm 0.83$ | $-22.40 \pm 1.63$ | $14.76 \pm 0.29$ | $21.53 \pm 0.32$ |
| $G_{\text{np}}$ | $-6.69 \pm 0.03$ | $-5.60 \pm 0.03$ | $-5.43 \pm 0.03$ | $-4.34 \pm 0.03$ |
| $\Delta H$ | $-48.50 \pm 0.36$ | $-32.82 \pm 0.38$ | $-34.96 \pm 0.27$ | $-26.37 \pm 0.30$ |
| $T\Delta S$ | $-23.81 \pm 0.67$ | $-20.12 \pm 0.38$ | $-19.98 \pm 0.42$ | $-19.86 \pm 0.12$ |
| $\Delta G_{\text{cal}}$ | $-24.69 \pm 0.52$ | $-12.70 \pm 0.38$ | $-14.98 \pm 0.34$ | $-6.51 \pm 0.21$ |

\*: The statistical error was estimated based on 0.5-5 ns MD simulation trajectory. 500 snapshots evenly extracted from the 0.5-5 ns MD trajectory of complex were used for MM/GBSA calculations and 50 snapshots for the entropy term calculations.

**Table 5.** Components of the Binding Free Energy (kcal/mol) Calculated by SIE Approach*.

| Energy term | nelfinavir | pitavastatin | perampanel | praziquantel |
| --- | --- | --- | --- | --- |
| $E_{\text{vdw}}$ | -58.36±0.32 | -42.77±0.30 | -44.33±0.25 | -34.92±0.32 |
| $E_C$ | -52.44±0.36 | 16.87±0.82 | 0.02±0.01 | -3.84±0.16 |
| $\Delta G_{\text{bind}}^R$ | 58.44±0.30 | -10.57±0.67 | 8.06±0.15 | 12.05±0.16 |
| $E_{\text{cavity}}$ | -9.93±0.05 | -7.79±0.04 | -8.19±0.04 | -6.66±0.06 |
| $\Delta G_{\text{cal}}$ | -9.42±0.04 | -7.53±0.04 | -7.55±0.03 | -6.39±0.04 |

\*: The binding free energy was calculated with the equation: $\Delta G_{\text{bind}} = 0.1048 * (E_C + E_{\text{vdw}} + \Delta G_{\text{bind}}^R + E_{\text{cavity}}) - 2.89$ . $E_{\text{cavity}} = \gamma \Delta \text{MSA}$ . The statistical error was estimated based on 0.5-5 ns MD simulation trajectory. 500 snapshots evenly extracted from the 0.5-5 ns MD trajectory were used.

### 3.3 Binding modes of nelfinavir against 2019-nCov M^pro^

As shown in Figure 2, the binding model of nelfinavir in its docking complex turned out to be very similar with that of the original ligand (TG-0205221)^25^ of 2GX4 (Figure 2a&b), which is an inhibitor of SARS M^pro^ with in vitro Ki of 0.053 μM and IC50 of 0.6 μM in the Vero-E6 cells.^25^ In its crystal structure, TG-0205221 is able to form hydrogen bonds with HIS-163, GLU-166 and GLN-189, and a very week hydrogen bond with PHE-140 (Figure 2c). Our docking results showed that three of the hydrogen bonds involving GLU-166 and GLN-189 maintained upon the binding of nelfinavir and 2019-nCov M^pro^, with additionally the possible formation of π-π stacking interaction with HIS-41 (Figure 2d). These observations further demonstrated that nelfinavir would interact with key residues of 2019-nCov M^pro^ in a similar way to that of the existing inhibitors against coronavirus M^pro^.

**Figure 2.**
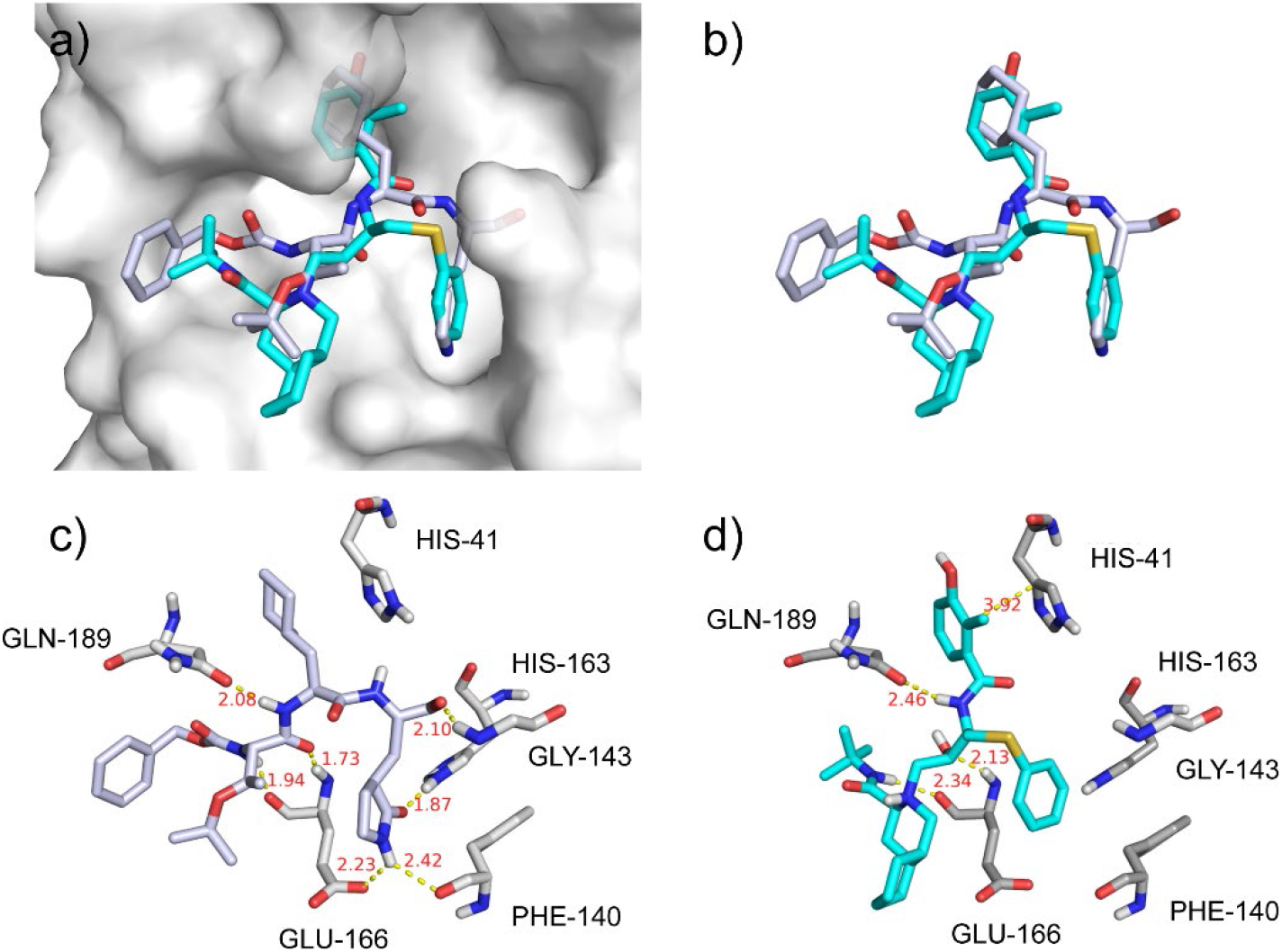
The binding model of nelfinavir against 2019-CoV M^pro^. (a) Binding models of the original ligand (TG-0205221, white) in 2GX4 and nelfinavir (cyan) in the 2019-nCov M^pro^ protein pocket (white surface); (b) Superposition of TG-0205221 (white) and nelfinavir (cyan) in their binding conformations; (c) Interactions between TG-0205221 and associated residues in the crystal structure (2GX4) of SARS M^pro^; (d) Interactions between nelfinavir and associated residues in the homology model of 2019-nCov M^pro^. The data in red is the interaction distance (Å).

## 4. Conclusions

2019-nCov caused more than 80 deaths in China as of 27 January 2020 and is a potential threat to the global health. However, there is no approved drug to treat the disease. 2019-nCov M^pro^ is a potential drug target to combat the virus, which shares 96% sequence similarity with the corresponding one in SARS. We built 11 homology models of 2019-nCov M^pro^ and docked 1903 approved small molecule drugs to the 2GTB model. Based on the docking score and the 3D similarity of binding mode to 39 known M^pro^ binders, 15 drugs were selected for further evaluation. The 15 drugs were then docked to all the 11 homology models, leading to 4 drugs for binding energy calculations. Both MM/GBSA and SIE calculations voted for nelfinavir, a HIV-1 protease inhibitor to treat HIV. Therefore, we suggested that nelfinavir might be active against 2019-nCov M^pro^. In addition, pitavastatin, perampanel, and praziquantel might also have moderate activities against 2019-nCov.

## Acknowledgments

This work was supported by the National Key R&D Program of China (2017YFB0202601). The simulations were partially run at TianHe 1 supercomputer in Tianjin. We thank Prof. Tingjun Hou in Zhejiang University, China for helpful discussions on the MM/GBSA calculations.

## References

1. Zhu, N.; Zhang, D.; Wang, W.; Li, X.; Yang, B.; Song, J.; Zhao, X.; Huang, B.; Shi, W.; Lu, R.; Niu, P.; Zhan, F.; Ma, X.; Wang, D.; Xu, W.; Wu, G.; Gao, G. F.; Tan, W., A Novel Coronavirus from Patients with Pneumonia in China, 2019. The New England journal of medicine 2020.

2. Lu, H.; Stratton, C. W.; Tang, Y.-W., Outbreak of Pneumonia of Unknown Etiology in Wuhan China: the Mystery and the Miracle. J. Med. Virol. 2020.

3. Hui, D. S.; I Azhar, E.; Madani, T. A.; Ntoumi, F.; Kock, R.; Dar, O.; Ippolito, G.; McHugh, T. D.; Memish, Z. A.; Drosten, C.; Zumla, A.; Petersen, E., The continuing 2019-nCoV epidemic threat of novel coronaviruses to global health - The latest 2019 novel coronavirus outbreak in Wuhan, China. International journal of infectious diseases: IJID: official publication of the International Society for Infectious Diseases 2020, 91, 264–266.

4. Chen, Y.; Liu, Q.; Guo, D., Coronaviruses: genome structure, replication, and pathogenesis. J. Med. Virol. 2020.

5. Jared S., M.; Tyler, L.; Shiqing, X.; Wenshe, L., Learning from the Past: Possible Urgent Prevention and Treatment Options for Severe Acute Respiratory Infections Caused by 2019-nCoV. 2020.

6. Burley, S. K.; Berman, H. M.; Bhikadiya, C.; Bi, C.; Chen, L.; Di Costanzo, L.; Christie, C.; Dalenberg, K.; Duarte, J. M.; Dutta, S.; Feng, Z.; Ghosh, S.; Goodsell, D. S.; Green, R. K.; Guranović, V.; Guzenko, D.; Hudson, B. P.; Kalro, T.; Liang, Y.; Lowe, R.; Namkoong, H.; Peisach, E.; Periskova, I.; Prlić, A.; Randle, C.; Rose, A.; Rose, P.; Sala, R.; Sekharan, M.; Shao, C.; Tan, L.; Tao, Y.-P.; Valasatava, Y.; Voigt, M.; Westbrook, J.; Woo, J.; Yang, H.; Young, J.; Zhuravleva, M.; Zardecki, C., RCSB Protein Data Bank: biological macromolecular structures enabling research and education in fundamental biology, biomedicine, biotechnology and energy. Nucleic Acids Res. 2018, 47 (D1), D464–D474.

7. Schrodinger, LLC, The PyMOL Molecular Graphics System, Version 2.4. 2019.

8. Waterhouse, A.; Bertoni, M.; Bienert, S.; Studer, G.; Tauriello, G.; Gumienny, R.; Heer, F. T.; de Beer, T. A P.; Rempfer, C.; Bordoli, L.; Lepore, R.; Schwede, T., SWISS-MODEL: homology modelling of protein structures and complexes. Nucleic Acids Res. 2018, 46 (W1), W296–W303.

9. Wishart, D. S.; Feunang, Y. D.; Guo, A. C.; Lo, E. J.; Marcu, A.; Grant, J. R.; Sajed, T.; Johnson, D.; Li, C.; Sayeeda, Z.; Assempour, N.; Iynkkaran, I.; Liu, Y.; Maciejewski, A.; Gale, N.; Wilson, A.; Chin, L.; Cummings, R.; Le, D.; Pon, A.; Knox, C.; Wilson, M., DrugBank 5.0: a major update to the DrugBank database for 2018. Nucleic Acids Res. 2018, 46 (D1), D1074–D1082.

10. Morris, G. M.; Huey, R.; Lindstrom, W.; Sanner, M. F.; Belew, R. K.; Goodsell, D. S.; Olson, A. J., AutoDock4 and AutoDockTools4: Automated docking with selective receptor flexibility. J. Comput. Chem. 2009, 30 (16), 2785–2791.

11. Koes, D. R.; Baumgartner, M. P.; Camacho, C. J., Lessons Learned in Empirical Scoring with smina from the CSAR 2011 Benchmarking Exercise. J. Chem. Inf. Model. 2013, 53 (8), 1893–1904.

12. Trott, O.; Olson, A. J., AutoDock Vina: Improving the speed and accuracy of docking with a new scoring function, efficient optimization, and multithreading. J. Comput. Chem. 2010, 31 (2), 455–461.

13. Dolinsky, T. J.; Nielsen, J. E.; McCammon, J. A.; Baker, N. A., PDB2PQR: an automated pipeline for the setup of Poisson–Boltzmann electrostatics calculations. Nucleic Acids Res. 2004, 32 (suppl_2), W665–W667.

14. Vaz de Lima, L. A.; Nascimento, A. S., MolShaCS: a free and open source tool for ligand similarity identification based on Gaussian descriptors. Eur. J. Med. Chem. 2013, 59, 296–303.

15. Wang, J.; Wolf, R. M.; Caldwell, J. W.; Kollman, P. A.; Case, D. A., Development and testing of a general amber force field. J. Comput. Chem. 2004, 25 (9), 1157–74.

16. Duan, Y.; Wu, C.; Chowdhury, S.; Lee, M. C.; Xiong, G.; Zhang, W.; Yang, R.; Cieplak, P.; Luo, R.; Lee, T.; Caldwell, J.; Wang, J.; Kollman, P., A point-charge force field for molecular mechanics simulations of proteins based on condensed-phase quantum mechanical calculations. J. Comput. Chem. 2003, 24 (16), 1999–2012.

17. Kollman, P. A.; Massova, I.; Reyes, C.; Kuhn, B.; Huo, S.; Chong, L.; Lee, M.; Lee, T.; Duan, Y.; Wang, W.; Donini, O.; Cieplak, P.; Srinivasan, J.; Case, D. A.; Cheatham, T. E., 3rd, Calculating structures and free energies of complex molecules: combining molecular mechanics and continuum models. Acc. Chem. Res. 2000, 33 (12), 889–97.

18. Srinivasan, J.; Cheatham, T. E.; Cieplak, P.; Kollman, P. A.; Case, D. A., Continuum Solvent Studies of the Stability of DNA, RNA, and Phosphoramidate-DNA Helices. J. Am. Chem. Soc. 1998, 120 (37), 9401–9409.

19. Naïm, M.; Bhat, S.; Rankin, K. N.; Dennis, S.; Chowdhury, S. F.; Siddiqi, I.; Drabik, P.; Sulea, T.; Bayly, C. I.; Jakalian, A.; Purisima, E. O., Solvated Interaction Energy (SIE) for Scoring Protein-Ligand Binding Affinities. 1. Exploring the Parameter Space. J. Chem. Inf. Model. 2007, 47 (1), 122–133.

20. Onufriev, A.; Bashford, D.; Case, D. A., Exploring protein native states and large-scale conformational changes with a modified generalized born model. Proteins 2004, 55 (2), 383–94.

21. Yang, H.; Xie, W.; Xue, X.; Yang, K.; Ma, J.; Liang, W.; Zhao, Q.; Zhou, Z.; Pei, D.; Ziebuhr, J.; Hilgenfeld, R.; Yuen, K. Y.; Wong, L.; Gao, G.; Chen, S.; Chen, Z.; Ma, D.; Bartlam, M.; Rao, Z., Design of wide-spectrum inhibitors targeting coronavirus main proteases. PLoS Biol 2005, 3 (10), e324.

22. Lee, T. W.; Cherney, M. M.; Huitema, C.; Liu, J.; James, K. E.; Powers, J. C.; Eltis, L. D.; James, M. N., Crystal structures of the main peptidase from the SARS coronavirus inhibited by a substrate-like aza-peptide epoxide. J Mol Biol 2005, 353 (5), 1137–51.

23. Ghosh, A. K.; Xi, K.; Ratia, K.; Santarsiero, B. D.; Fu, W.; Harcourt, B. H.; Rota, P. A.; Baker, S. C.; Johnson, M. E.; Mesecar, A. D., Design and synthesis of peptidomimetic severe acute respiratory syndrome chymotrypsin-like protease inhibitors. J Med Chem 2005, 48 (22), 6767–71.

24. Lee, T. W.; Cherney, M. M.; Liu, J.; James, K. E.; Powers, J. C.; Eltis, L. D.; James, M. N., Crystal structures reveal an induced-fit binding of a substrate-like Aza-peptide epoxide to SARS coronavirus main peptidase. J Mol Biol 2007, 366 (3), 916–32.

25. Yang, S.; Chen, S. J.; Hsu, M. F.; Wu, J. D.; Tseng, C. T.; Liu, Y. F.; Chen, H. C.; Kuo, C. W.; Wu, C. S.; Chang, L. W.; Chen, W. C.; Liao, S. Y.; Chang, T. Y.; Hung, H. H.; Shr, H. L.; Liu, C. Y.; Huang, Y. A.; Chang, L. Y.; Hsu, J. C.; Peters, C. J.; Wang, A. H.; Hsu, M. C., Synthesis, crystal structure, structure-activity relationships, and antiviral activity of a potent SARS coronavirus 3CL protease inhibitor. J. Med. Chem. 2006, 49 (16), 4971–80.

26. Lu, I. L.; Mahindroo, N.; Liang, P. H.; Peng, Y. H.; Kuo, C. J.; Tsai, K. C.; Hsieh, H. P.; Chao, Y. S.; Wu, S. Y., Structure-based drug design and structural biology study of novel nonpeptide inhibitors of severe acute respiratory syndrome coronavirus main protease. J Med Chem 2006, 49 (17), 5154–61.

27. Goetz, D. H.; Choe, Y.; Hansell, E.; Chen, Y. T.; McDowell, M.; Jonsson, C. B.; Roush, W. R.; McKerrow, J.; Craik, C. S., Substrate specificity profiling and identification of a new class of inhibitor for the major protease of the SARS coronavirus. Biochemistry 2007, 46 (30), 8744–52.

28. Ghosh, A. K.; Xi, K.; Grum-Tokars, V.; Xu, X.; Ratia, K.; Fu, W.; Houser, K. V.; Baker, S. C.; Johnson, M. E.; Mesecar, A. D., Structure-based design, synthesis, and biological evaluation of peptidomimetic SARS-CoV 3CLpro inhibitors. Bioorg Med Chem Lett 2007, 17 (21), 5876–80.

29. Verschueren, K. H.; Pumpor, K.; Anemuller, S.; Chen, S.; Mesters, J. R.; Hilgenfeld, R., A structural view of the inactivation of the SARS coronavirus main proteinase by benzotriazole esters. Chem Biol 2008, 15 (6), 597–606.

30. Lee, C. C.; Kuo, C. J.; Ko, T. P.; Hsu, M. F.; Tsui, Y. C.; Chang, S. C.; Yang, S.; Chen, S. J.; Chen, H. C.; Hsu, M. C.; Shih, S. R.; Liang, P. H.; Wang, A. H., Structural basis of inhibition specificities of 3C and 3C-like proteases by zinc-coordinating and peptidomimetic compounds. J Biol Chem 2009, 284 (12), 7646–55.

31. Zhu, L.; George, S.; Schmidt, M. F.; Al-Gharabli, S. I.; Rademann, J.; Hilgenfeld, R., Peptide aldehyde inhibitors challenge the substrate specificity of the SARS-coronavirus main protease. Antiviral Res 2011, 92 (2), 204–12.

32. Jacobs, J.; Grum-Tokars, V.; Zhou, Y.; Turlington, M.; Saldanha, S. A.; Chase, P.; Eggler, A.; Dawson, E. S.; Baez-Santos, Y. M.; Tomar, S.; Mielech, A. M.; Baker, S. C.; Lindsley, C. W.; Hodder, P.; Mesecar, A.; Stauffer, S. R., Discovery, synthesis, and structure-based optimization of a series of N-(tert-butyl)-2-(N-arylamido)-2-(pyridin-3-yl) acetamides (ML188) as potent noncovalent small molecule inhibitors of the severe acute respiratory syndrome coronavirus (SARS-CoV) 3CL protease. J Med Chem 2013, 56 (2), 534–46.

33. Kim, Y.; Lovell, S.; Tiew, K. C.; Mandadapu, S. R.; Alliston, K. R.; Battaile, K. P.; Groutas, W. C.; Chang, K. O., Broad-spectrum antivirals against 3C or 3C-like proteases of picornaviruses, noroviruses, and coronaviruses. J Virol 2012, 86 (21), 11754–62.

34. Turlington, M.; Chun, A.; Tomar, S.; Eggler, A.; Grum-Tokars, V.; Jacobs, J.; Daniels, J. S.; Dawson, E.; Saldanha, A.; Chase, P.; Baez-Santos, Y. M.; Lindsley, C. W.; Hodder, P.; Mesecar, A. D.; Stauffer, S. R., Discovery of N-(benzo[1,2,3]triazol-1-yl)-N-(benzyl)acetamido)phenyl) carboxamides as severe acute respiratory syndrome coronavirus (SARS-CoV) 3CLpro inhibitors: identification of ML300 and noncovalent nanomolar inhibitors with an induced-fit binding. Bioorg Med Chem Lett 2013, 23 (22), 6172–7.

35. Shimamoto, Y.; Hattori, Y.; Kobayashi, K.; Teruya, K.; Sanjoh, A.; Nakagawa, A.; Yamashita, E.; Akaji, K., Fused-ring structure of decahydroisoquinolin as a novel scaffold for SARS 3CL protease inhibitors. Bioorg Med Chem 2015, 23 (4), 876–90.

36. Tomar, S.; Johnston, M. L.; St John, S. E.; Osswald, H. L.; Nyalapatla, P. R.; Paul, L. N.; Ghosh, A. K.; Denison, M. R.; Mesecar, A. D., Ligand-induced Dimerization of Middle East Respiratory Syndrome (MERS) Coronavirus nsp5 Protease (3CLpro): IMPLICATIONS FOR nsp5 REGULATION AND THE DEVELOPMENT OF ANTIVIRALS. J Biol Chem 2015, 290 (32), 19403–22.

37. St John, S. E.; Tomar, S.; Stauffer, S. R.; Mesecar, A. D., Targeting zoonotic viruses: Structure-based inhibition of the 3C-like protease from bat coronavirus HKU4--The likely reservoir host to the human coronavirus that causes Middle East Respiratory Syndrome (MERS). Bioorg Med Chem 2015, 23 (17), 6036–48.

38. St John, S. E.; Therkelsen, M. D.; Nyalapatla, P. R.; Osswald, H. L.; Ghosh, A. K.; Mesecar, A. D., X-ray structure and inhibition of the feline infectious peritonitis virus 3C-like protease: Structural implications for drug design. Bioorg Med Chem Lett 2015, 25 (22), 5072–7.

39. Galasiti Kankanamalage, A. C.; Kim, Y.; Damalanka, V. C.; Rathnayake, A. D.; Fehr, A. R.; Mehzabeen, N.; Battaile, K. P.; Lovell, S.; Lushington, G. H.; Perlman, S.; Chang, K. O.; Groutas, W. C., Structure-guided design of potent and permeable inhibitors of MERS coronavirus 3CL protease that utilize a piperidine moiety as a novel design element. Eur J Med Chem 2018, 150, 334–346.

